# Obesity Disrupts Pituitary UPR Leading to NAFLD

**DOI:** 10.1101/2022.07.06.498610

**Authors:** Qingwen Qian, Mark Li, Zeyuan Zhang, Shannon Davis, Kamal Rahmouni, Andrew W. Norris, Huojun Cao, Wen-Xing Ding, Ling Yang

## Abstract

Obesity is the major risk factor for nonalcoholic fatty liver disease (NAFLD), for which effective cures are lacking. Despite the notion that obesity is associated with aberrant levels and action of pituitary hormones that are essential for maintaining hepatic metabolic and inflammatory states, the intrinsic pituitary endocrine abnormalities and their systemic consequences are incompletely defined. By characterizing the impact of diet-induced obesity (DIO) on the pituitary whole tissue and single cell transcriptome, we demonstrated that obesity disrupts pituitary endoplasmic reticulum (ER) homeostasis by suppressing the inositol-requiring enzyme-α (IRE1α)-mediated adaptive unfolded protein response (UPR). We further showed that defective pituitary UPR by IRE1α-deletion in the anterior pituitary strikingly augmented obesity-associated systemic metabolic abnormalities, particularly the NAFLD-associated pathologies. Conversely, enhancing the adaptive UPR in the anterior pituitary, by genetic gain-of-function of spliced X-box binding protein 1 (sXBP1), ameliorated the systemic and hepatic metabolic defects observed in mice with pituitary IRE1α deletion. Intriguingly, disruption of the UPR in the pituitary resulted in impaired hepatic UPR, which was in part due to a defective thyroid hormone receptor (THR)-mediated activation of hepatic *Xbp1*. In contrast, activation of the hepatic THR signaling improved obesity-associated glucose intolerance and attenuated the impaired hepatic ER homeostasis in anterior pituitary-IRE1α deficient mice. Together, our study provides the first insight into disruption of endocrine signaling-mediated inter-organ UPR communication drives obesity-associated hepatic pathologies. Unraveling these connections might uncover new therapeutic targets for NAFLD and other obesity-associated diseases.

## INTRODUCTION

Obesity increases the risk for development of a wide range of metabolic diseases including insulin resistance, type 2 diabetes and nonalcoholic fatty liver disease (NAFLD)^1^. It is estimated that NAFLD occurs in 70-80% of individuals with obesity^2,3^, posing a substantial public health burden. The pituitary gland synthesizes and secretes hormones essential for maintaining hepatic lipid and glucose metabolic homeostasis, as well as its inflammatory state^4,5^. Obesity disrupts both the level and action of pituitary hormones in rodents and humans^6–8^. Of note, the prevalence of NAFLD is increased in patients with hypopituitarism^9^, growth hormone (GH) deficiency^10^ and hypothyroidism^11,12^, contributing to premature mortality. It is well recognized that pituitary hormone secretion is controlled by the hypothalamic neuroendocrine input^13,14^, and obesity disrupts hypothalamic neuroendocrine function, which has been implicated in the consequential impaired releasing of regulatory pituitary hormones^15,16^. In parallel, accumulating evidence demonstrates an interplay of obesity-associated hypothalamic cellular dysfunction and hepatic glucose and lipid metabolism defects in the context of pathogenesis of NAFLD^17–19^. However, despite the notion that the pituitary gland coordinates both hypothalamic and peripheral inputs, it is unknown to what extent this hypothalamus-to-liver regulation is pituitary-dependent, and whether an intrinsic pituitary endocrine dysfunction directly leads to NAFLD progression.

Up to 70% of the pituitary gland mass is comprised of professional secretory cells^20^, which require an intact function of the endoplasmic reticulum (ER) to support their normal endocrine action. Perturbation of ER homeostasis causes ER stress, which activates the unfolded protein response (UPR) in order to restore normal ER function^21^. It is established that failing to initiate this adaptive response to mitigate ER stress leads to abnormalities in other endocrine secretory cells such as impaired insulin secretion in β-cells of the pancreas^22–24^. The kinetics of pituitary hormone production are tightly controlled by nutritional flux, which is a physiological inducer of ER stress^21,22^. Thus, pituitary function may be particularly sensitive to ER stress. Indeed, a recent study in a cell culture system indicated that ER stress contributes to pituitary adenoma progression^23^, and ER stress markers are present in the pituitary of rats fed a high-sucrose diet and high-fat diet^24,25^. However, the pituitary UPR profile and its direct functional significance in the context of NAFLD have not been characterized.

In the liver, failure to activate the adaptive arm of the UPR to restore ER homeostasis is a major factor in the pathogenesis of steatosis, insulin resistance and inflammation in NAFLD and its advanced form, non-alcoholic steatohepatitis (NASH)^26–28^. Although the mechanisms by which the protective hepatic UPR is lost are largely unknown, alleviation of ER stress by chemical or molecular chaperones improves metabolic control and insulin sensitivity in both obese mice and humans^29–33^. Systemically, it has been implicated that the hepatic UPR senses endocrine inputs to modulate hepatic cellular and metabolic programs. For example, it is shown that the hepatic UPR is acutely activated by elevation of insulin upon feeding^34^. In addition, catabolic activation of the hepatic adaptive UPR by glucagon and epinephrine signaling improves the disrupted ER function in the setting of NAFLD^35,36^. However, it is largely unknown whether and to what extent aberrant pituitary hormone signaling contributes to hepatic UPR disruption in the setting of obesity.

Organisms adapt and respond to environmental challenges by coordinating stress responses across cells, tissues and organs. In the liver, hepatocyte ER stress can be propagated between neighboring cells^37^. Although cell-autonomous regulation of the UPR has been extensively studied, the regulatory mechanisms of the UPR that communicate stress across organs/tissues throughout the organism are incompletely understood, especially in the context of obesity-associated metabolic diseases. Here, we demonstrated that obesity desensitized the UPR in the pituitary. We further provide the first insights into a thyroid hormone-mediated pituitary-to-liver UPR interplay as well as its functional relevance in the context of NAFLD.

## RESULTS

### Obesity disrupts the pituitary UPR response

To gain a broad understanding of whether and to what extent obesity alters nutritional flux-induced responses in pituitary, we performed an RNA-seq analysis using the pituitary from lean mice (fed normal chow diet; RD) and diet-induced obese mice (DIO, fed a high fat diet, HFD), under fasting and refed conditions. In lean mice, nutritional flux markedly altered the pituitary transcriptome (SFig. 1A & SFig. 1B), while this effect was largely blunted in DIO mice (SFig. 1C & SFig. 1D). Gene ontology (GO) biological process analysis revealed that inflammation and lipid metabolism were the top-enriched pathways that were induced in the pituitary by obesity (Fig. 1A). Notably, the UPR signaling pathway was not only the most significantly enriched signature following refeeding in lean mice (SFig. 1B), but also the most significantly downregulated in DIO mice (Fig. 1A).

**Figure 1.**
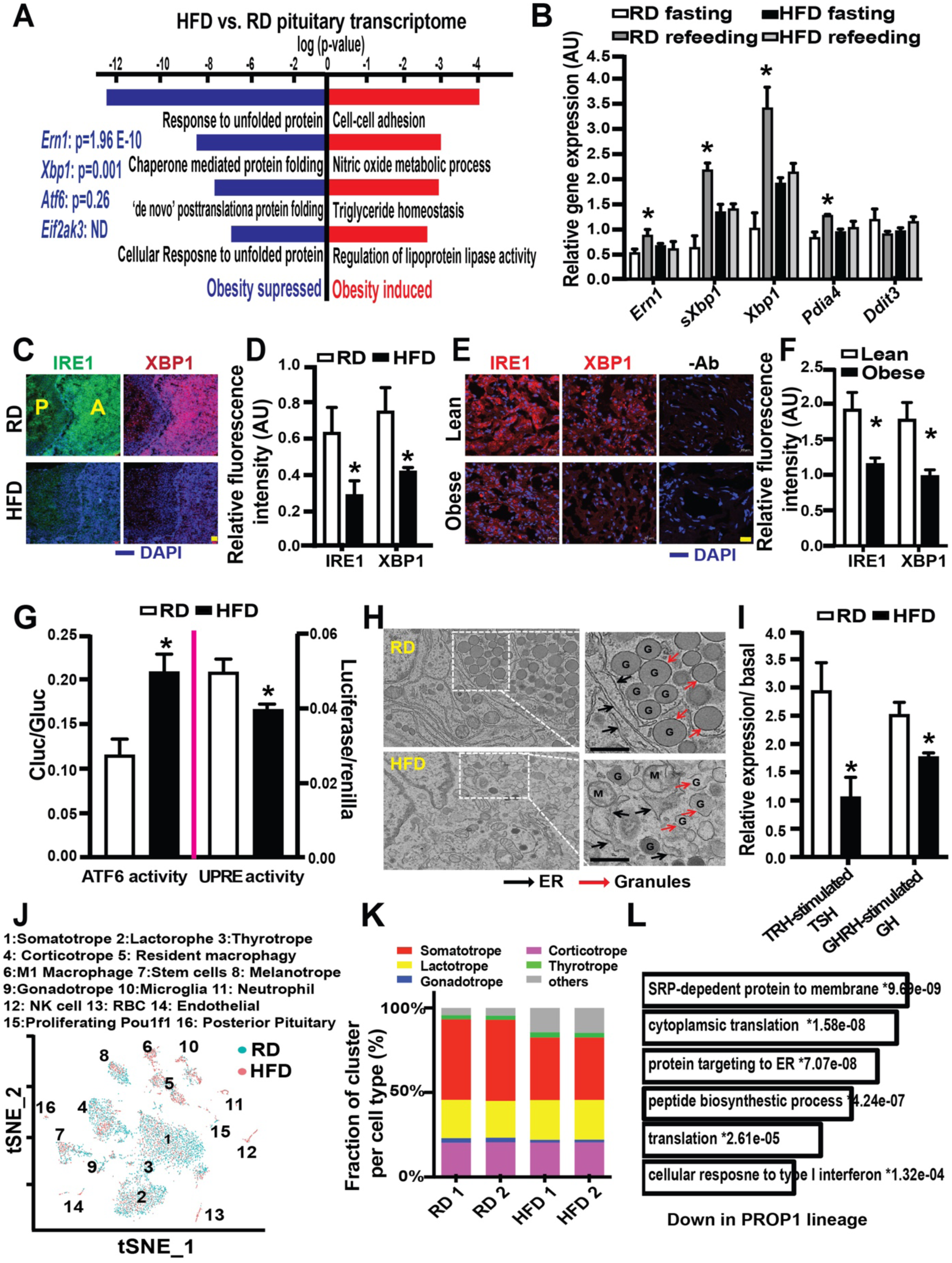
Obesity impairs the IRE1α-XBP1 arm of pituitary UPR. **A.** RNA-Seq data analyses (Gene Ontology [GO] Biological process) of significantly enriched pathways in the pituitary of refed male mice (4 hrs following a 16-hrs fasting) on a HFD (16 wks on HFD) vs. mice on a RD, n= 3 mice/group. Significantly altered genes are listed on the left of the panel. **B.** Levels of mRNAs encoding genes of interest (normalized to *Hprt1*), and **C-D.** Representative images (C) and quantification (D) of tested proteins in the pituitary from RD and HFD (16 wks on HFD) male mice with a 4-hrs refeeding following a 16-hrs fasting, n= 4 mice/group. Scale bar: 50 μm. **E-F.** Representative images (E) and quantification (F) of tested proteins in human pituitary (n= 2 individuals/group). Scale bar: 20 μm. **G.** ATFLD-cluc secretion (left), and UPRE-luc activity (right) from isolated primary anterior pituitary cells from C57BL/6J mice fed with RD and HFD for 16 wks (n= 5-7mice/group). Data were normalized to Gluc secretion or Renilla luciferase respectively. **H.** Representative TEM micrographs of pituitary from RD or HFD mice. **I.** TSH and GH levels in primary anterior pituitary cells from mice in (C), followed by stimulation with TRH (500 ng/ml, 1 hr) or GHRH (8 μM, 1 hr). **J-K.** T-distributed stochastic neighbor embedding (t-SNE) plots of pituitary cell clusters (J), and percent contribution of anterior endocrine cell clusters in scRNA-Seq analysis (K) for pituitary of refed male mice on a RD or a HFD (16 wks on HFD). **L.** GO analysis of suppressed DEGs in PROP1 linage of pituitary cells from HFD mice in (J). Data in (B, D, F, G and I) are presented as means ± SEM. * indicates refeeding effects in (B), dietary effects in (D, G and I), and compared to the lean group in (F), as determined by ANOVA in (B) and Student’s t-test in (D, F, G and I), p<0.05. AU: arbitrary unit.

In response to ER stress, inositol-requiring enzyme-α (IRE1α) processes the X-box binding protein 1 (XBP1) mRNA to a spliced form (sXBP1), which then is translated and acts as a transcription factor to upregulate a subset of UPR target genes involved in protein folding and quality control, thereby attenuating stress levels^38^. We found that obesity significantly suppressed pituitary *Ern1* (gene encoding IRE1) and s*Xbp1* expression in response to feeding (Fig. 1A&B). Immunohistochemical analyses further supported our transcriptomic results, showing that in the anterior pituitary, obesity significantly suppressed IRE1α and XBP1 (Fig. 1C&D) but not other UPR branches such as protein kinase RNA-like ER kinase (PERK) or its downstream regulator C/EBP homologous protein (CHOP) (SFig. 1E). Notably, we observed a decrease in pituitary levels of IRE1α and XBP1 in obese humans (Fig. 1E&F). To directly measure the effect of obesity on ER function, we isolated primary anterior pituitary cells from lean and DIO mice, and monitored ER homeostasis using an activating transcription factor 6 luminal domain (ATF6LD) *cypridina noctiluca (Cluc)* secretion reporter. In this assay, cells with abundant ER chaperone activity retain ATF6LD-*Cluc* in the ER, whereas reduced chaperone availability results in a release of this chimeric protein and luciferase secretion into the medium^39^. In these same cells, we also monitored IRE1α-mediated sXBP1 activation using a UPR response element (UPRE, sXBP1-specific DNA binding site) luciferase reporter^40^. As shown in Fig. 1G, obesity induced ER stress (elevated ATF6-cluc in the medium), but impaired the adaptive UPR (suppressed UPRE activity) in pituitary cells.

We next utilized transmission electron microscopy (TEM) to visualize the ultracellular architecture of the pituitary from lean and obese mice. As shown in Fig. 1H, obesity completely disrupted granule morphology and ER structure in the hormone-producing cells in the pituitary. Given that the pituitary hormone production is regulated by the hypothalamic hormonal input^13,14^, it is possible that this obesity-associated maladaptive pituitary UPR might be due to lower protein synthesis demand caused by insufficient hypothalamic hormonal release. Therefore, we next assessed thyrotropin-releasing hormone (TRH)-stimulated thyroid-stimulating hormone (TSH), and growth hormone-releasing hormone (GHRH)-stimulated GH release from freshly isolated anterior pituitary of lean and obese mice. In obese mice, the effect of hypothalamic releasing hormones (TRH and GHRH) on TSH and GH secretion was severely blunted (Fig. 1I), indicating an intrinsic defect in the pituitary hormone secretion in obese mice. This aberrant endocrine function of the pituitary was not related to lower number of residency cells or apoptosis in pituitary from obese mice (SFig. 1F), as determined by flow cytometry analysis.

To delineate the impact of obesity on the population-specific UPR landscape, we further performed a single cell RNA sequencing (scRNA-Seq) analysis of the pituitary from lean and obese mice. Hormones produced by the anterior pituitary govern systemic metabolic homeostasis and energy balance^20^. We found that obesity did not skew the general anterior pituitary cell composition in male mice, except for lowering the somatotrope signature and enriching other cell types such as immune cell population (e.g., macrophages) (Fig. 1J&K). Notably, obesity suppressed the UPR landscape in major anterior pituitary endocrine cell populations: thyrotrope (which secrete thyroid-stimulating hormone), lactotrophe (which secrete prolactin), and somatotrope (which secrete growth hormone [GH])^41,42^(Fig. 1L). These data demonstrated that obesity disrupts the intra-pituitary endocrine function, which is correlated with a maladaptive UPR.

### IRE1**α** is required for anterior pituitary function to maintain systemic energy and metabolic homeostasis

To investigate whether there is a direct impact of dysfunctional pituitary IRE1-XBP1 UPR branch on systemic metabolic homeostasis, we generated mice with IRE1α deletion in the anterior pituitary (IRE1^PKO^) by crossing IRE1^fl^ mice with Prophet of Pit 1 (*Prop 1*) Cre mice^43^. PROP1 is a member of the paired-like family of homeodomain transcription factors, which is required for the development of thyrotropes, lactotrophes, and somatotropes. These mice were also further bred with ROSA^mT/mG^ reporter mice^44^. The IRE1^PKO^ mice were viable and fertile, and as shown in Fig. 2A, *Prop1* Cre-mediated deletion of IRE1α occurred in the anterior but not in the posterior pituitary. Furthermore, scRNA-Seq analysis revealed that loss of IRE1α in male mice did not alter pituitary cell composition (Fig. 2B & SFig. 2A), but decreased the UPR transcriptome and s*Xbp1* expression in the pituitary PROP1 lineage cells as expected (SFig. 2B and Fig. 2C&D). This defect in the UPR signature was neither associated with a reduction of the pituitary hormone transcripts we measured nor in the number of anterior pituitary cells (SFig. 2C&D).

**Figure 2.**
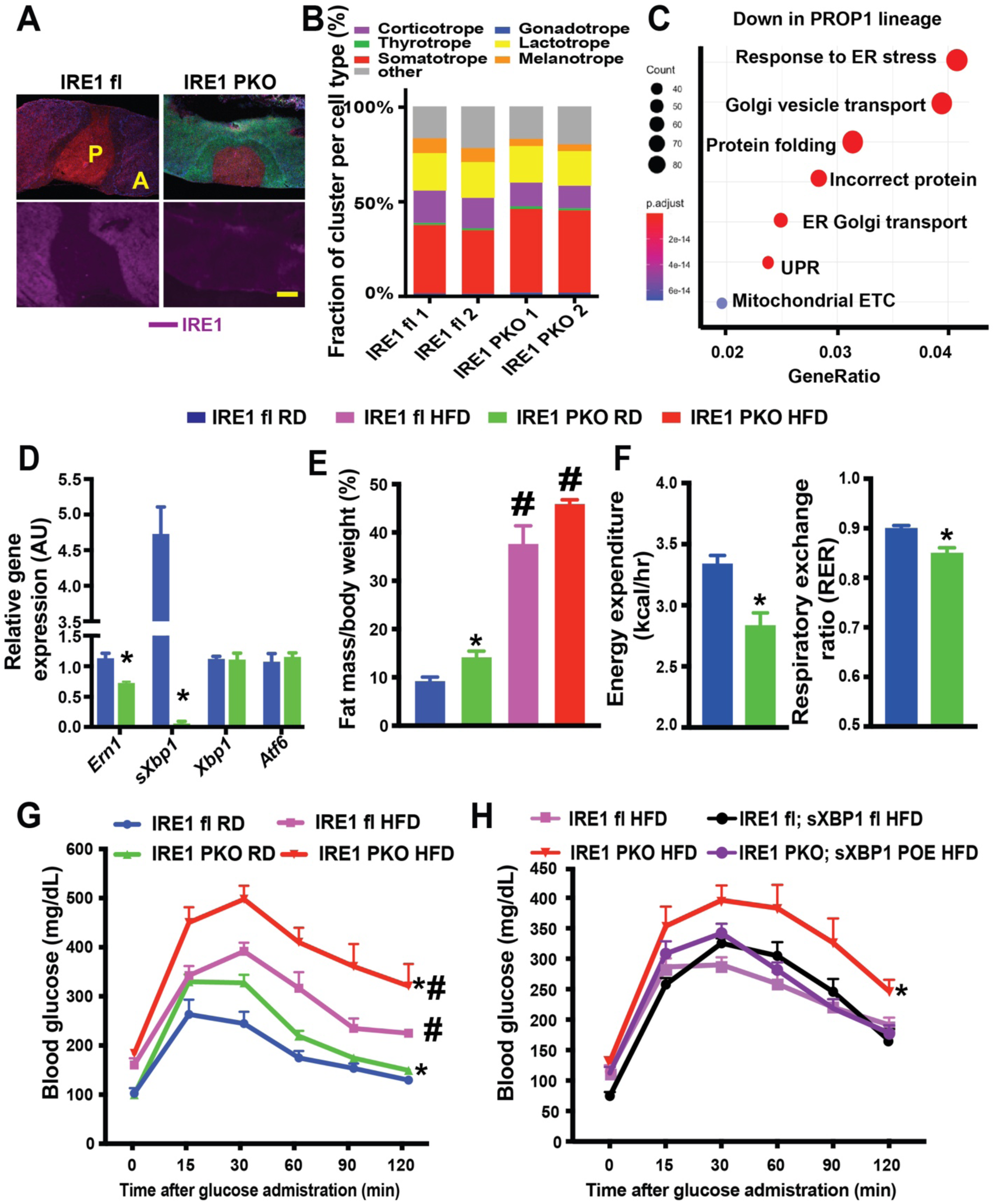
Anterior pituitary IRE1α-sXBP1 axis is necessary and sufficient to protect against obesity-associated metabolic dysfunction. **A.** Representative images of GFP, tdTomato and staining of IRE1α in the pituitary from IRE1^fl^ and IRE1^PKO^ mice. A: anterior pituitary, P: posterior pituitary. Expression of GFP indicates the deletion of IRE1 in pituitary. Scale bar: 200 μm. **B.** Percent contribution of pituitary endocrine cell clusters in scRNA-Seq analysis for IRE^fl^ and IRE1^PKO^ mice fed with RD. **C.** GO analysis of down-regulated DEGs in PROP1 linage from IRE1^PKO^ mice compared with IRE1^fl^ mice. **D.** Levels of mRNA encoding gene of interest in pituitary from IRE1^fl^ and IRE1^PKO^ mice (n= 3 mice/group) were determined by quantitative RT-PCR. Data were normalized to the expression of *Hprt1*. **E.** Body composition of IRE1^fl^ and IRE1^PKO^ mice fed a RD or a HFD for 16 wks (n= 5-6 mice/group) were measured by nuclear magnetic resonance. **F.** Whole body energy expenditure and respiratory exchange ratio (RER) in IRE1^fl^ and IRE1^PKO^ mice fed a RD (n= 5-6 mice/group). **G.** Glucose tolerance test (GTT) from IRE1^fl^ and IRE1^PKO^ mice fed a RD or a HFD for 16 wks (n= 8-10 mice/group). **I.** GTT from IRE1^fl^, IRE1^PKO^, IRE1^fl^; sXBP1^fl^, and IRE1^PKO^; sXBP1^POE^ mice fed mice fed a HFD for 12 wks (n= 6-8 mice/group). Data in (D-H) are presented as means ± SEM. * indicates genetic effects in (D-H) in mice on same diet, and # indicates dietary effects in same type of mice in (E and G); as determined by Student’s t-test in (D and F), by ANOVA in (E), and by ANOVA followed by post-hoc test of AUC in (G and H), p<0.05. AU: arbitrary unit.

Given that DIO is more pronounced in male mice than female mice^45^, we first assessed the effects of pituitary IRE1α deletion on systemic metabolism in male mice. IRE1α anterior pituitary deletion did not alter the weight of either lean or obese mice (SFig. 2E), but whole-body composition analysis revealed that IRE1^PKO^ mice had larger fat mass and reduced less lean mass compared to their controls (Fig. 2E and SFig. 2F). We then examined whether IRE1α in the anterior pituitary is involved in the regulation of systemic energy balance. Indirect mouse calorimetry analysis showed that IRE1^PKO^ lean mice had reduced energy expenditure (EE) and respiratory exchange ratio (RER) compared to controls (Fig. 2F), which was not associated with significant changes in food intake (SFig. 2G). Moreover, IRE1^PKO^ mice displayed impaired glucose tolerance under RD, which was worsened by HFD (Fig. 2G). On the other hand, we did not observe significant alteration in serum insulin levels between HFD groups (SFig. 2H). IRE1α possesses both a kinase and an RNase domain in the cytosolic portion of the protein^21^. Previously, it is demonstrated that XBP1 splicing signaling attenuates cellular and metabolic stres^38^, whereas persistent activation of IRE1α kinase suppresses insulin signaling in obesity^46^. To determine whether gain-of adaptive IRE1α signaling function in the pituitary could rescue the metabolic defects caused by disruption of pituitary UPR, we crossed IRE1^fl^ mice with mice conditionally overexpressing sXBP1 (sXBP1^fl^, OE)^47^. These mice were then bred with *Prop1*-Cre mice to overexpress sXBP1 and simultaneously delete IRE1α in the anterior pituitary (IRE1^PKO^; sXBP1^POE^). We found that restoration of pituitary sXBP1 in IRE1^PKO^ HFD mice improved glucose intolerance to the level observed in IRE1^fl^ controls (Fig. 2H & SFig. 2I). Together these data indicate that the pituitary IRE1-sXBP1 axis is necessary and sufficient for maintaining systemic metabolic homeostasis in health and obesity.

### Anterior pituitary IRE1**α** protects against NAFLD-associated liver pathologies and ER dysfunction

The pituitary hormone plays a critical role in modulating hepatic metabolic function^48^. To investigate the impact of the pituitary UPR dysfunction on the liver, we performed an RNA-seq analysis in livers from obese IRE1^fl^ and IRE1^PKO^ mice. Pathway analysis revealed that pituitary IRE1α deletion significantly enriched transcriptomic signatures of hepatic fibrosis and steatosis in DIO mice, but suppressed the hepatic ER transcriptome (Fig. 3A and SFig. 3A-C). Consistent with these transcriptional analyses, deletion of IRE1α in the pituitary increased hepatic steatosis, triglyceride (TG) content in both lean and obese mice (Fig. 3B&C). Obesity-induced NAFLD results in mild inflammation and fibrosis^49^. To further determine the extent to which the pituitary UPR dysfunction contributes to liver pathologies, we challenged mice with a high-fat, high cholesterol and high fructose diet (HFHCHFr) to induce mouse model of NASH^50^. As shown in Fig. 3D-F, deletion of IRE1α in the pituitary increased macrophage infiltration and fibrosis in livers of mice on both RD and HFHCHFr diet. Notably, the hepatic transcriptional signature in the IRE1^PKO^ HFD mice was comparable to that observed in our previous study of mice with hepatic deletion of *Xbp1*^47^(Fig. 3G), as well as an independently published transcriptome dataset of livers from mice with liver-specific XBP1^51^ (SFig. 3D). To determine whether disruption of the pituitary UPR affects hepatic XBP1 action, we assessed expression of *Xbp1* and its targets in livers from IRE1^PKO^ mice and its controls. Indeed, impairment of the pituitary UPR suppressed hepatic *sXbp1* and sXBP1 targeted genes (Fig. 3H), consistent with elevated expression of genes involved in lipogenesis and gluconeogenesis (SFig. 3E). Notably, restoration of sXBP1 in the anterior pituitary ameliorated steatosis and immune cell infiltration in the liver of obese IRE1^PKO^ mice (Fig. 3I-K). Together, our data indicate that pituitary UPR dysfunction contributes to NAFLD progression and UPR dysfunction in the liver.

**Figure 3.**
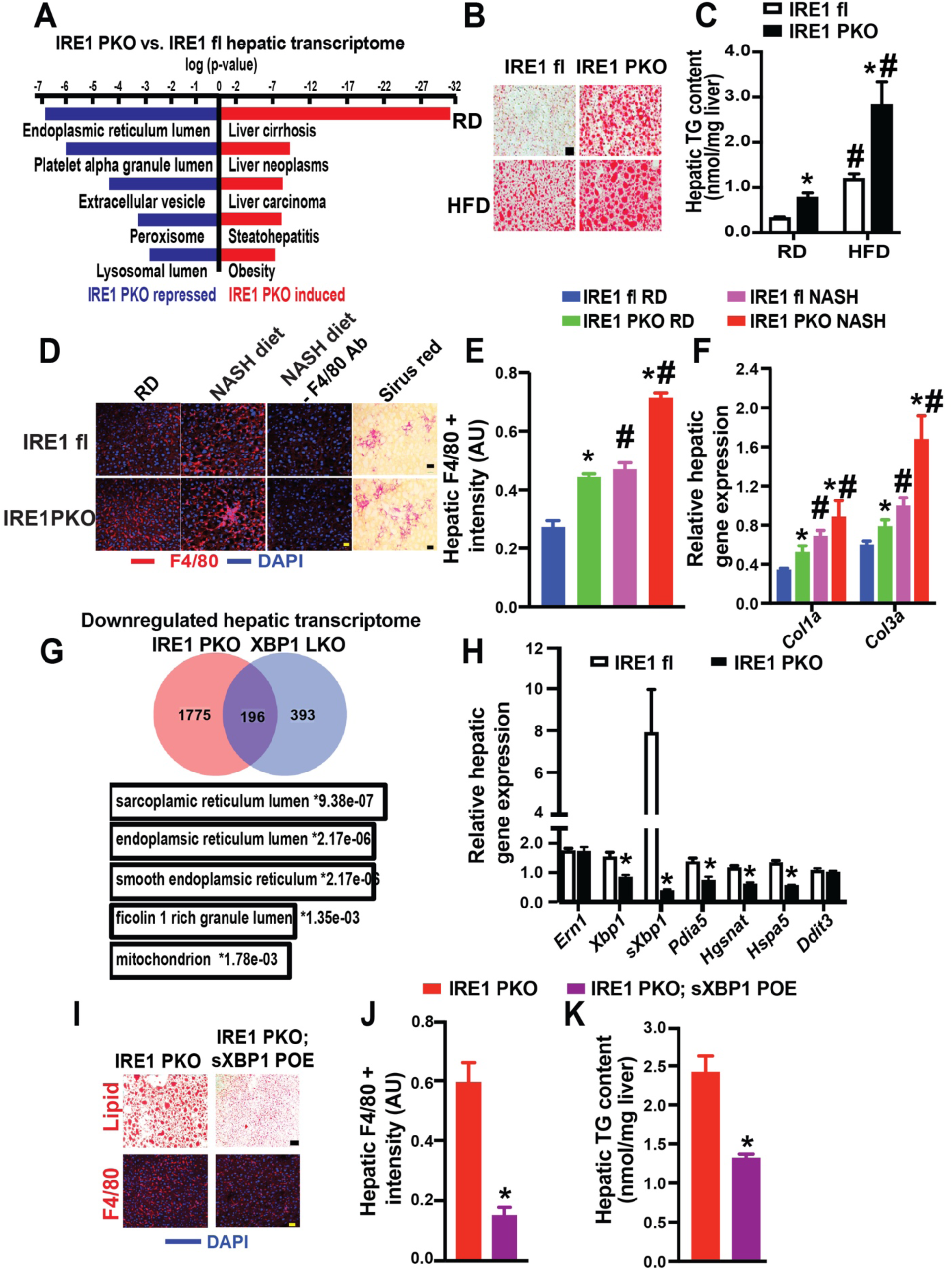
IRE1α pituitary deficiency contributes to NAFLD progression. **A.** RNA-Seq data analyses (GO biological process and DisGeNET) of livers from male IRE1^fl^ and IRE1^PKO^ mice on a HFD for 20 wks (n= 3 mice/group). **B.** Representative images of Oil Red O staining (scale bar: 100μm) and **C.** Hepatic triglyceride (TG) levels in mice (n= 3 mice/group) in (B). **D.** Representative images of F4/80 (scale bar: 20μm), and Sirius Red staining (scale bar: 100 μm), **E.** Quantification of F4/80 staining, and **F**. Levels of mRNA encoding genes of interest (data were normalized with *Hprt1*) in livers from IRE1^fl^ and IRE1^PKO^ mice fed a RD or NASH diet for 33 wks (n= 3 mice/group). **G.** RNA-Seq data analyses of DEGs in livers from IRE1^fl^, IRE1^PKO^, XBP1^LKO^ and XBP1^fl^ mice fed a HFD for 16 wks mice (n= 3 mice/group). Top panel: the common down-regulated DEGs signatures (IRE1^PKO^ v.s. IRE1^fl^, XBP1^LKO^ v.s. XBP1^fl^) and Bottom panel: GO analysis of common down-regulated DEGs. **H.** Levels of mRNAs encoding genes of interest in livers from mice in (A), as assessed by quantitative RT-PCR. Data were normalized to *Hprt1*. n= 3 mice/group. **I.** Representative images of Oil Red O staining (scale bar: 100 μm) and F4/80 (scale bar: 20 μm), **J.** Quantification of F4/80 staining in (I), and **K.** Hepatic TG levels in mice in IRE1^PKO^ and IRE1^PKO^; sXBP1^POE^ mice fed mice fed a HFD for 12 wks (n= 3 mice/group). Data are presented as means ± SEM in (C, E, F, H, J and K). * Indicates genetic effects in mice on same diet, # indicates dietary effects in same type of mice; as determined by Student’s t-test in (H, J and K), by ANOVA in (C, E and F) followed by post-hoc test, p<0.05. AU: arbitrary unit.

### Thyroid hormone signal links pituitary UPR action on the hepatic UPR

Pituitary-derived hormones play a critical role in modulating hepatic lipid and glucose metabolism^4,5^. Epidemiological studies have shown an inverse relationship between serum levels of TH (a key hormone for maintaining hepatic lipid homeostasis) and the incidence of NAFLD, and patients with NASH have double the prevalence of hypothyroidism^11,57^. TH synthesis and secretion is modulated by the hypothalamus-pituitary-thyroid axis, involving hypothalamic secretion of TRH, which acts on thyrotropes in the anterior pituitary to produce TSH, and subsequent TSH-stimulated TH release from the thyroid to maintain systemic euthyroid range. Our pathway analysis of scRNA-Seq revealed that the *Ern1* (gene encodes IRE1)-*Xbp1 axis* was the most enriched downregulated UPR signature in thyrotrope cells in obese mice compared to lean mice (Fig. 4A). These transcriptomic features were concomitant with reduced circulating T3, as well as reduced intrahepatic T3 in lean and obese IRE1^PKO^ mice (Fig. 4B-C).

**Figure 4.**
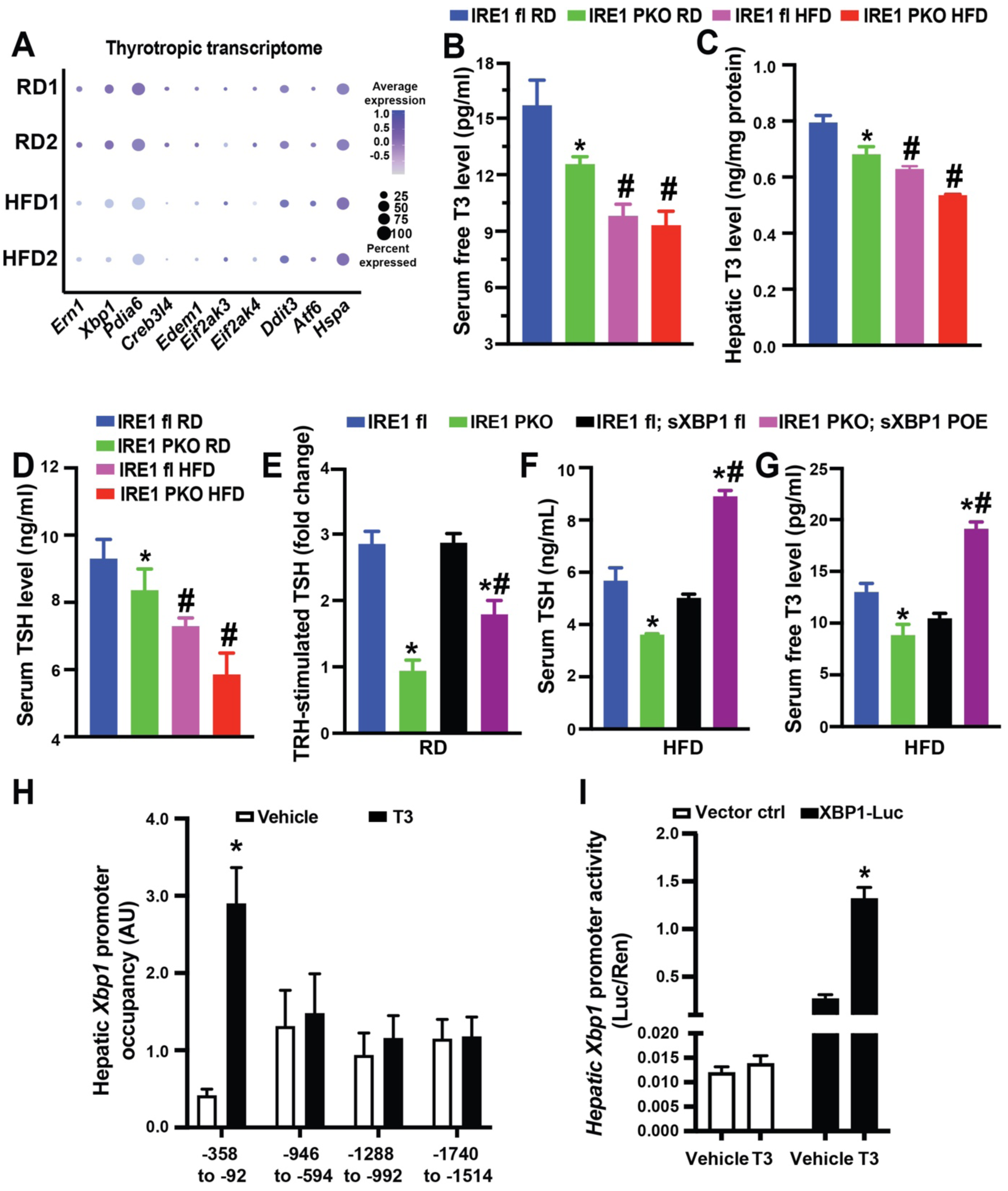
Loss of pituitary IRE1α impairs action of pituitary–thyroid axis on hepatic *Xbp1*. **A.** Dot plots of thyrotrope-UPR markers in pituitary scRNA-Seq of mice on a RD or a HFD (16 wks on HFD). **B.** Serum free T3 levels, **C.** Hepatic T3 levels, and **D**. Serum TSH levels in IRE1^fl^ and IRE1^PKO^ mice (on diet for 20 wks) by ELISA. n= 5-6 mice/group. **E.** TRH-stimulated TSH levels in anterior pituitaries from IRE1^fl^, IRE1^PKO^, IRE1^fl^; sXBP1^fl^ and IRE1^PKO^; sXBP1^POE^ mice fed a RD for 20 wks (n=3 mice/group), followed stimulation with TRH (500 ng/ml, 1 hr). **F-G.** Serum levels of hormones of interest in IRE1^fl^, IRE1^PKO^, IRE1^fl^; sXBP1^fl^ and IRE1^PKO^; sXBP1^POE^ mice fed on a HFD for 20 wks. n= 3 mice/group**. H.** Occupancy of *Xbp1* promoter regions by TH receptor β (THRB) in primary hepatocytes isolated from C57BL/6J (WT) mice treated with T3 (500 nM, 8 hrs). n= 3 mice/group. **I**. Activity of *Xbp1* promoter (−350 to −198) in primary hepatocytes from WT mice following transfection with the indicated constructs (CT: PGL4.10) in the presence or absence of T3 (500 nM) for 8 hrs. The data were normalized to Renilla luciferase. n= 3 experimental replicates. Data are presented as means ± SEM in (B-I). * Indicates genetic effects in mice on same diet in (B-G), and treatment effects in same type of cell in (H and I); # indicates dietary effects in same type of mice in (B-G); as determined Student’s t-test in (H and I), and ANOVA followed by post-hoc test in (B-G), p<0.05. AU: arbitrary unit.

Circulating levels of TSH are maintained by a balance between central TSH production and peripheral TSH clearance, and only being significantly dropped at the onset of severe and critical illness or in response to some medication^52,53^. We found that although TSH levels were slightly elevated (potentially due to a compensatory response) in young IRE1^PKO^ mice compared to littermate controls, they were decreased in adult IRE1^PKO^ mice and were further downregulated by exposure to HFD feeding (SFig. 4B & Fig. 4D). To determine the impact of IRE1α deficiency on central TSH production, we measured TRH-stimulated TSH release from anterior pituitary cells freshly isolated from lean IRE1^PKO^ mice and littermate controls. Pituitary IRE1 deficiency suppressed TRH-induced TSH production from the anterior pituitary in lean mice (Fig. 4E). This defective stimulated pituitary hormone secretion was also detected in freshly isolated anterior pituitary cells with *sXbp1* suppression (SFig. 4C&D). Importantly, restoration of anterior pituitary sXBP1 rescued TRH-stimulated TSH release from anterior pituitary cells with IRE1α deletion (Fig. 4E) and improved circulating TSH and T3 levels in mice on a HFD (Fig. 4F&G). Together, these data indicate that disruption of the pituitary UPR attenuates central and peripheral pituitary TH hormonal regulation.

The active form of TH, T3, controls gene expression in target tissues by binding to its cognate nuclear receptors (THRα and THRβ)^54,55^, which bind to TH response elements (TRE) in the promoter regions of target genes^54^. In an *in-silico* analysis, we found several potential TRE sites on the promoter region of *Xbp1* (Fig. 4H). To establish the direct regulation of hepatic *Xbp1* by TH, we assessed THRβ binding to the promoter of *Xbp1* by performing ChIP analysis using primary hepatocytes from WT RD mice using an anti-THRβ antibody. As shown in Fig. 4H, THRβ occupied the promoter of *Xbp1* at the −358 to −92 site in the presence of T3. Next, to test whether TH activates *Xbp1*, we generated an *Xbp1* luciferase reporter construct that contains the THR binding sites within the *Xbp1* promoter region (−350 to −198). As shown in Fig. 4I, T3 increased *Xbp1* promoter activity in primary hepatocytes. These findings suggested that TH signaling directly activates *Xbp1*, thereby regulating the hepatic UPR.

### The effect of pituitary UPR action on NAFLD is thyroid hormone signaling-dependent

Epidemiological studies have shown an inverse relationship between serum levels of TH and the incidence of NAFLD^11,56,57^. To determine whether TH can rescue the hepatic UPR defect in IRE1^PKO^ mice, we treated obese IRE1^PKO^ mice and its littermate controls with a liver-specific THR agonist, MGL-3196, which is currently in a phase III clinical trial^58^. As shown in Fig. 5A-C, activation of hepatic THR signaling improved obesity-associated glucose intolerance and hepatic steatosis. Notably, MGL-3196 treatment significantly increased THR occupancy on the *Xbp1* promoter in the liver (Fig. 5D).

**Figure 5.**
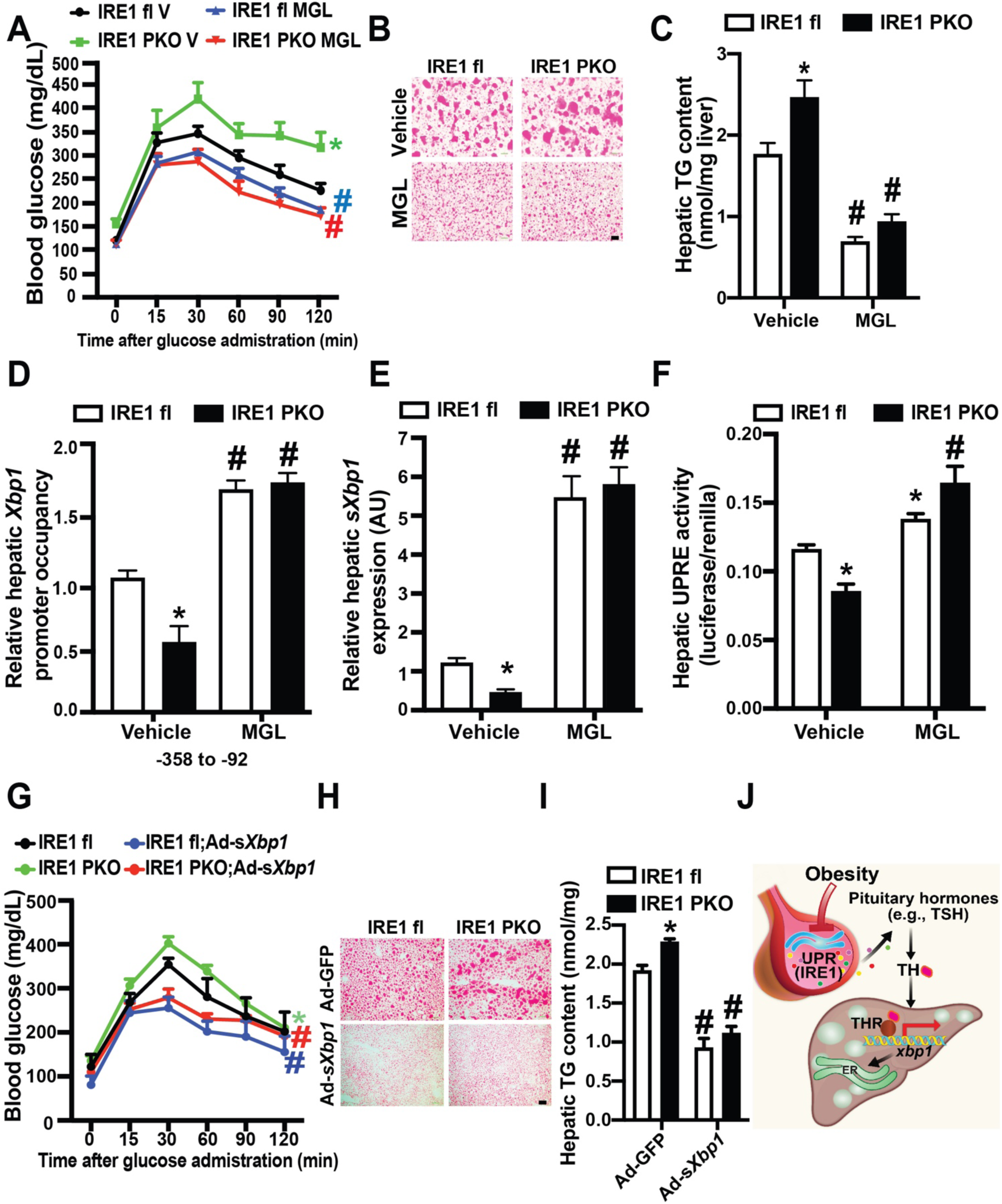
Activation of hepatic TH–*Xbp1* axis ameliorates defective pituitary UPR-associated metabolic dysfunction in the context of obesity. **A.** GTT in mice fed a HFD for 16 wks, following administration of MGL-3196 at a dose of 1mg/kg/day for 3 wks. V: vehicle. n= 6-11 mice/group **B.** Representative images of Oil Red O staining, and **C.** Hepatic TG levels in the liver from mice in (A). n= 3 mice/group **D.** Occupancy of *Xbp1* promoter regions by THRβ in livers of HFD mice treated with MGL-3196 or vehicle as in (A) by ChIP assay. For all ChIP assays, data were first normalized to 2% input and then to IgG. **E.** Levels of *sXbp1* transcript (data were normalized to *Hprt*) in livers of mice in (A), as assessed by qRT-PCR. n= 3 mice/group. **F**. UPRE-luc activities from primary hepatocytes isolated from IRE1^fl^ and IRE1^PKO^ mice on a RD in the presence or absence of MGL-3196 (22 μM, 8 hrs). n= 3 mice/group, data were normalized to Renilla luciferase. **G.** GTT of HFD mice (16 wks on diet) with adeno-*sXbp1* (LOE) or control viral transduction (8X10^10^ vp/mouse, 2 wks). n= 6-12 mice/group. **H.** Representative images of Oil Red O staining and **I.** Hepatic TG contents in liver from mice in (G). n= 3-6 mice/group. **J.** Model depicting pituitary UPR-mediated hepatic UPR in the context of NAFLD. All data are presented as means ± SEM. *Indicates genetic effects in mice with same treatments, and # indicates treatment effects in same type of mice as determined by ANOVA in (C-F and I) and ANOVA of AUC in (A and G) followed by post-hoc test, p<0.05. AU: arbitrary unit.

Obesity-associated ER dysfunction is characterized by impaired adaptive sXBP1 signaling in the liver^27,59^. To assess whether amelioration of hepatic TH action improves the adaptive hepatic UPR, we examined sXBP1 action in primary hepatocytes from IRE1^PKO^ mice and littermate controls in the presence or absence of MGL-3196. As shown in Fig. 5E&F, pituitary IRE1α deficiency in HFD mice decreased hepatic *sXbp1* expression as well as hepatocyte sXBP1 activity, which was reversed by treatment with MGL-3196. Finally, to determine whether the protective role of the pituitary UPR on hepatic function is mediated by sXBP1 in the liver, we transduced adeno-sXBP1 into livers of IRE1^PKO^ mice and littermate controls on a HFD (SFig. 4E). Hepatic sXBP1 overexpression alleviated systemic glucose homeostasis (Fig. 5G) and hepatic steatosis in both IRE1^fl^ and IRE1^PKO^ mice (Fig. 5H&I, and SFig. 4E). Furthermore, the effect of insulin on suppression of glucose production was significantly attenuated in hepatocytes from IRE1^PKO^ mice compared to controls, but this was ameliorated by overexpression of *sXbp1* (SFig. 4F). Collectively, these results indicate that TH serves as a modulator that links the pituitary IRE1-sXBP1 axis to the hepatic sXBP1 signaling, pointing to dysfunction in TH signaling as a potential physiological cause of the impaired hepatic UPR in the context of NAFLD (Fig. 5G).

## DISSCUSSION

The hypothalamic-pituitary (H-P) axis senses systemic inputs and responds to these cues by modulating multiple pituitary hormones to maintain peripheral tissue homeostasis. An extensive body of studies has primarily focused on the relevance of the hypothalamic inputs to peripheral metabolic homeostasis in the context of obesity. So far, however, there has been little direct evidence of whether and to what extent obesity promotes an intrinsic pituitary defect, as well as whether the resulting endocrine defect directly drives NAFLD progression. In our study, we demonstrated that obesity disrupts the ability of the adaptive pituitary UPR to mitigate ER stress induced by nutritional flux, and the resultant endocrine dysfunction contributes to hepatic metabolic abnormalities by impairing the hepatic UPR (Fig. 5J).

We found that obesity suppresses the pituitary IRE1-XBP1 branch of the UPR, the most conserved component of adaptive UPR signaling. IRE1α possesses both a kinase and an RNase domain in the cytosolic portion^21^, with two outputs of IRE1 RNase activity, namely XBP1 splicing and IRE1-dependent RNA decay (RIDD)^60,61^. We showed that overexpressing sXBP1 while simultaneously deleting IRE1α in the anterior pituitary alleviated systemic and hepatic metabolic abnormalities in IRE1^PKO^ mice, indicating that the endocrine regulatory role of IRE1α in the anterior pituitary is in part through mediating the RNase action. Future work evaluating the contribution of IRE1α-mediated RIDD and IRE1α kinase activity in pituitary in the context of obesity is needed.

Our transcriptomic and IHC analyses did not reveal a significant alteration of the other UPR branches, ATF6 and PERK, in the pituitary of obese mice. This observation is in line with a recent study demonstrating that XBP1 branch is the main physiological UPR pathway that responds to the high secretory demand in cultured pituitary cells^62^. Selective disruption of the UPR by obesity has been shown in other peripheral organs such as the liver and pancreas, wherein PERK signaling is sustained but IRE1-sXBP1 signaling is downregulated^27,63,64^. However, the mechanism underlying this selective disruption is unknown. One potential factor is inflammation. It has been shown that hypothalamic inflammation contributes to systemic energy and metabolic imbalance in the context of obesity^15,65^ in part attribute to proinflammatory cytokines-mediated inhibition of hypothalamic release of regulatory pituitary hormones^15,16^. Indeed, our scRNA-Seq analysis in the pituitary revealed that obesity elevates proinflammatory signatures (*eg. Tnf* and *Nfkb1* levels) in both pituitary endocrine and immune cell populations (unpublished data). Therefore, future studies will focus on potential contributions of obesity-associated inflammation on the pituitary UPR defect.

At the cellular level, ER stress and dysregulation of the UPR are considered as pathological “hits” of NAFLD progression ^26,66,67^. At the physiological level, the hepatic UPR is regulated by multiple endocrine inputs. For example, the hepatic UPR is acutely activated in response to the elevation in insulin^34^. A recent study also demonstrated activation of the hepatic adaptive UPR by the catabolic signals glucagon and epinephrine signaling^35^, which could improve the disrupted ER function in the setting of NAFLD^36^. We previously showed that XBP1 is regulated during lactation, and responds to prolactin to alter lipogenic gene expression in the adipose tissue^68^. A very recent study showed that GH activates XBP1 in rat liver in a sexually dimorphic manner via ERK and the C/EBPβ pathway^69^. Here, we revealed that hepatic *Xbp1* is directly activated by TH signaling. Together, these observations support the notion that disruption of pituitary endocrine inputs is one of the key factors that leads to obesity-associated malfunction of ER in peripheral tissues.

TH plays a key role in the regulation of homeostasis and energy maintenance, and its dysfunction is linked to diverse diseases including NAFLD^56,57^ and NASH^11^. The systemic “thyrostat” is controlled centrally and peripherally by multiple complex hormonal feedback loops and enzymatic processes. It is suggested that a change in circulating levels of TSH that parallels with changes in T3/T4 indicates a defect originating in the anterior pituitary. In contrast, an inverse relationship between changes of TSH and T3/T4 indicates thyroid gland dysfunction as a primary cause^70–72^. We found that circulating TSH levels were slightly elevated in in young IRE1^PKO^ mice compared to littermate controls (SFig. 4A). We postulate that this is in part due to a compensatory response to lower T3, which is commonly observed in patients with subclinical hypothyroidism^73,74^. However, long-term HFD feeding and IRE1α pituitary deletion lowered serum TSH and T3 levels in older adult mice with age, indicating pituitary endocrine deficiency as a primary cause. Indeed, we demonstrated that both DIO and pituitary IRE1-deficiency suppressed central TSH production (Fig. 1H and Fig. 4E). In the future, it will be informative to compare this finding in mice to the variation in TSH levels in obese humans.

Currently, the causal relationships between primary/secondary hypopituitarism and primary/secondary hypothyroidism in the context of NAFLD are largely unclear. Our study provides the first insight into the causal role of defective pituitary UPR on TH abnormalities as its related to NAFLD progression, reflecting a state of primary hypopituitarism-to-secondary hypothyroidism. Although it is unclear whether obesity is a mild central hypothyroidism in humans, it is well documented in rodent studies that obesity disrupts central thyroid responses^7,75,76^. It is possible that the pituitary UPR plays a role in setting the central “thyrostat” checkpoint or that there are inputs from other hypothalamic neurons to the pituitary. We found a direct transcriptional regulatory axis of THRβ-*Xbp1* in the liver. Hence, it is plausible that obesity-associated central thyroid desensitization results in an impaired THRb-mediated activation of *Xbp1* in the pituitary, which in turn results in endocrine defects. While our current work demonstrates that disruption of the pituitary-to-liver TH signaling serves as a potential physiological cause of the impaired hepatic UPR in the context of NAFLD, the direct impact of pituitary UPR on the peripheral thyrostat, such as thyroid gland function, requires further characterization.

The systemic propagation of organelle stress responses has been evidenced in both non-mammalian and mammalian systems^77^. In *Caenorhabditis elegans* activation of sXBP1 in the nervous system evokes the UPR in intestinal cells and promotes lysosomal biogenesis^78^. In the liver, intercellular hepatocyte communication propagates obesity-associated ER stress signals via Cx43-mediated signal transduction^37^. In mice, cold-induced autophagy in proopiomelanocortin (POMC) neurons induces autophagy in the brown adipose tissue and the liver^79^. Furthermore, selective overexpression of sXBP1 in the POMC neurons leads to increased *Xbp1* mRNA splicing in the liver^17^. However, it’s currently unclear how POMC neurons communicate their local organelle stress response to the liver. Our study provided the first evidence that pituitary hormones can serve as modulators that directly link the UPR across organs *in vivo*, as well as its functional relevance in the context of NAFLD. Together, given the strong evidence that pituitary hormones serve as cell non-autonomous signals that activate the hepatic UPR, targeting pituitary UPR function may be a feasible strategy to prevent interorgan propagation of ER dysfunction and thus represent a novel approach to alleviate obesity-associated metabolic diseases including NAFLD.

## ACKNOWLEDGMENTS

We thank Dr. Takao Iwawaki at the Gunma University (Japan) and Dr. Gökhan S. Hotamisligil (Harvard T. H. Chan School of Public Health) for providing the IRE1^fl^ mouse. We also thank Dr. Hotamisligil for providing the sXBP1^fl^ mice, and the UPRE-luc and ATF6-cluc constructs. We are grateful to Drs. Dale Abel (University of Iowa) and Julien Sebag (University of Iowa) for scientific discussions and insights. We also thank Dr. Sara C. Sebag for helping editing this manuscript, and all Yang laboratory members for supporting this project.

## AUTHOR CONTRIBUTIONS

L.Y. and Q.Q. designed the study and wrote the manuscript. Q.Q., M.L., Z.Z., H.C. and W.X.D. performed the experiments. S.D., K.R., and A.W.N. provided critical reagents and scientific suggestions on the manuscript. L.Y. conceived and supervised the study, and is supported by NIH R01 DK108835-01A1.

## MATERIALS AND METHODS

### Mouse models, diet and treatments

All the animal care and experimental procedures were performed with approval from the University of Iowa’s Institutional Animal Care and Use Committee. All mice were bred and maintained under specific pathogen free conditions, and temperature was maintained at 22°C. Mice were kept on a 12 hrs light-dark cycle and fed with RD (Teklad global diet, 7913). Mice used in generating the DIO model were placed on a 60% kCal high-fat diet (Research Diets, D12492) immediately after weaning, at 3 wks of age. Mice used for NASH model were fed with high-fat, high cholesterol and high fructose (HFHCHFr; Research diet, D09100310). C57BL/6J mice were purchased from Jackson Laboratory. s*Xbp1*^fl^ mice (Hprt^tm1(fl-STOP-fl-sXbp1)Hota^) were generated by Dr. Gökhan S. Hotamisligil (Harvard T. H. Chan School of Public Health). IRE1^fl^ mice were from Dr. Takao Iwawaki at the Gunma University (Japan). The *Prop1-cre* mice were generated, and transferred from the laboratory of Dr. Shannon Davis at the University of South Carolina. For MGL-3196 interventional studies, mice were fed with 60% HFD for the indicated time and randomly assigned to the treatment and control groups. Mice were then treated with 1 mg/kg MGL-3196 (Cayman chemical, Cat No. 23845) or 0.9% NaCl (vehicle) via intraperitoneal injection (i.p.) daily for 3 wks. Adenovirus-*GFP* and adenovirus-s*Xbp1* were delivered *via* retro-orbital injection at a titer of 1X10^11^ ifu/mouse^27,80,81^. After 6 hrs food withdrawal, tissues were harvested, frozen in liquid nitrogen, and kept at −80°C until processing. At the end of the experiment, mice were scarified, and organs and serum were collected for the further analysis.

### Primary hepatocytes culture, treatments and analyses

Mouse liver was perfused with collagenase type X (Wako, 039-17864), and the cells were washed with hepatocyte wash medium (Thermo Fisher Scientific, Cat No. 17704024)^80^. After purification by Percoll (GE Healthcare Life Sciences, Cat No. 17089101) density gradient separation, cells were resuspended in William’s E medium (Thermo Fisher Scientific, Cat No. 12551032) with 5% FBS (Gibco, Cat No. 26140079), 10 nM dexamethasone (Sigma-Aldrich, Cat No. D1756), and 20 nM INS (insulin, Sigma-Aldrich, Cat No. I5500). Hepatocytes were then seeded on COL1A1/collagen I (Corning, Cat No. 354236)-coated plates at a final density of 3.5 × 10^4^ cells/cm^2^. Cells were cultured with fresh medium and transduced with the indicated adenoviruses after 4 hrs of attachment or transfected with indicated constructs after 16 hrs of attachment. The concentration of T3 was 500nM, MGl-3196 was 22 μM.

#### Glucose production in isolated primary hepatocytes

Primary hepatocytes were cultured with serum free William’s E medium for 16 hrs, then treated with 20 nM insulin (Sigma-Aldrich, Cat No. I5500), 10 μM forskolin (Sigma, Cat No. F6886) for 6 hrs in phenol red-free William’s E medium (Lonza, Cat No. BE02-019F). Glucose level in supernatant was measured using 2,2′-Azino-bis(3-ethylbenzothiazoline-6-sulfonic acid)- diammonium salt (ABTS)-linked glucose oxidase-peroxidase assay^82^. Briefly, 50μl of medium was used in each reaction and the absorbance was measured at 420nm on a microplate reader (SpectraMax, Molecular Devices, San Jose, CA). The protein concentration was determined to normalize the values of glucose production.

#### Adenovirus transduction and transient transfection

For adenovirus transduction, 4 MOI of adenovirus was added to isolated primary hepatocytes. To measure ER homeostasis, isolated primary hepatocytes were seeded to pre-coated rat tail collagen I (Corning, Cat No. 354236) 24-well plates. After overnight incubation, cells were transfected with the 0.6 µg/well ATF6LD-Cluc-Gluc^39^ or UPRE-luciferase reporter^40^ with 0.15 µg/well Renilla luciferase vector (Promega, Cat No. E2261) using polyethylenimine (PEI, Polysciences INC, Cat No. 23966). At 48 hrs post-transfection, the activities of Cypridina noctiluca (Clu) luciferase in medium were measured using 1 mM Cypridina cypridina Luciferin (Prolume, Cat No. #305), and the Gaussia luciferase (Gluc) was used as a secretion control measured using 10 mM native coelenterazine (CTZ) (Prolume, Cat No. #303). In the UPRE reporter assay, firefly luciferase and Renilla luciferase were measured using the Dual-Glo Luciferase Assay (Promega, Cat No. E2920). The luminescence was read on a microplate reader (SpectraMax, Molecular Devices, San Jose, CA).

### Primary anterior pituitary cell culture, treatments and analyses

Isolation of anterior pituitary cells was performed as previously described^83^ with minor modification. Briefly, freshly isolated anterior pituitary was rinsed with Hank’s balanced salt solution (HBSS) buffer (Gibco, Cat No. 14175095), then minced in digestion buffer containing 2 mg/ml Collagenase, Type 2 (Worthington Biochemical Corporation, Cat No. LS004176), 3% BSA and in HBSS with 25 mM HEPES (Gibco, Cat No. 15630080) at 37°C for 1 hr with gentle shaking. Dispersed cells were then washed with HBSS (free of calcium and magnesium) at 100×g for 3 min at 4°C, resuspended in DMEM/F12 medium (Gibco, Cat No. 11320033) containing 15% fetal bovine serum (Gibco, Cat No. 16140071), and seeded to 24-well cell culture plates (Corning, Cat No. 3524). Cells were allowed to attach for 1–2 days at 37°C before treatment.

#### ER homeostasis analyses and promoter activity assay

Anterior pituitary cells were transfected with the 0.6 µg/well ATF6LD-Cluc-Gluc^39^ or UPRE-luciferase reporter^40^ with 0.15 µg/well Renilla luciferase vector (Promega, E2261) using Lipofectamine 3000 reagent (Thermo Fisher, L3000015, Waltham, MA) in Opti-MEM medium (Thermo Fisher Scientific, 31985-062, Waltham, MA). The *Xbp1* promoter containing putative THRβ binding site was synthesized from −350bp to −198bp (5’- CTC TAG TGT TTT CTT TTA AAG TTA TTA ATT TAT ATC AAT TAA TTT ATA TGA AAT AGG GTT TGA TCT ATA GCC CCA GCT GGC CTA GCT AGA CCT CCC TAT GTA GCC CAG GCT GC TCT CAA ACT GGC AAT AAA GTG CTG AGA TTA CAA GTG AGT A -3’) with restricted digestion enzymes site of KpnI and HindIII, and cloned into pGL4.15 luciferase reporter vector (Promega, E6701). At 48 hrs post-transfection, the activities of Clu, Gluc, firefly and Renilla luciferases were measured as described above, and the luminescence was read on a microplate reader (SpectraMax, Molecular Devices, San Jose, CA). *Flow cytometry analysis*

Single pituitary suspended cells from experimental mice were fixed by 4% paraformaldehyde solution and permeabilized by Intracellular Staining Perm Wash Buffer (Biolegned, Cat No. 421002). Resuspended cells were incubated with antibodies against GH (Santa Cruz, Cat No. sc-374266, 1:500), TSH (Santa Cruz, Cat No. sc-365801, 1:500), cleaved Caspase-3 (Cell Signaling, Cat No. 9664, 1:500), and Lamin A/C (Cell Signaling, Cat No. 4777, 1:500), followed by 1 hr incubation with fluorescent-conjugated secondary antibodies at RT for 1hr. Cell population was analyzed using a Cytek Northern Light (Cytek Biosciences). The number of events was stopped at 50,000 counts. Data collected from the experiments were analyzed using FlowJo 10.0.

#### Stimulated-pituitary hormone secretion

TRH-stimulated TSH secretion was performed as the described by Fleckman et al^84^ and GHRH-stimulated GH secretion was performed as the described by Lecoq et al^85^. with modifications. Briefly, anterior pituitaries were minced by blade and washed with 3% BSA in Krebs-Ringer Bicarbonate Buffer (KRB; 120 mM NaCl, 4.8 mM KCl, 2.5 mM CaCl_2_, 1.2 mM MgCl_2_, 10 mM HEPES, and 24 mM NaHCO_3_) for 4 hrs, followed with treatment of GHRH (8 μM; Sigma, Cat No. SCP0160) or TRH (500 ng/ml, Sigma, Cat No. P1319) in KRB buffer with 3% BSA for additional 1 hr. Supernatants from cells at the before and at the end of stimulation were collected, and cells were lysed at the end of treatment. Levels of TSH and GH were determined by enzyme-linked immunoassay (ELISA) using commercially available ELISA kits (TSH ELISA kit, G-Biosciences, Cat No. IT6045; GH ELISA kit, Crystal Chem, Cat No. 80587) respectively according to manuals. The stimulated hormone secretion was normalized to the basal levels.

### Quantitative real-time RT-PCR

Total RNA from liver and pituitary was isolated using the Trizol reagent (Invitrogen, Cat No. 15596026) and quantified with Nanodrop. cDNA was generated using an iScript cDNA synthesis kit (BioRad, Cat No. 1708891). Quantitative real-time PCR analysis was performed using BioRad CFX96 System. The primers used in the mouse study were: *Atf6* forward 5’- TCG TGT TCT TCA ACT CAG CAC-3’, *Atf6* reverse 5’- TGG AGT CAG TCC ATG TTC TGT-3’; *Col1a* forward 5’- CCA CGT CTC ACC ATT GGG G -3’, *Col1a* reverse 5’ GGC CCG GGA AGT CAC TGT -3’; *Col3a* forward 5’- CTG TAA CAT GGA AAC TGG GGA AA-3’, *Col3a* reverse 5’ CCA TAG CTG AAC TGA AAA CCA CC -3’; *Ddit3* forward 5’- CCA CCA CAC CTG AAA GCA GAA -3’, *Ddit3* reverse 5’- AGG TGA AAG GCA GGG ACT CA -3’; *Edem* forward 5’- AAG CCC TCT GGA ACT TGC G -3’, *Edem* reverse 5’- AAC CCA ATG GCC TGT CTG G -3’; *Ern1* forward 5’- ATG GCT ACC ATT ATC CTG AGC A -3’, *Ern1* reverse 5’- TCC TGG GTA AGG TCT CCG TG -3’; *Fasn* forward 5’- AGA GAT CCC GAG ACG CTT CT -3’, *Fasn* reverse 5’- GCC TGG TAG GCA TTC TGT AGT -3’; *Fshb* forward 5’- GCC ATA GCT GTG AAT TGA CCA-3’, *Fshb* reverse 5’- AGA TCC CTA GTG TAG CAG TAG C -3’; *G6Pase* forward 5’- CGA CTC GCT ATC TCC AAG TGA -3’, *G6Pase* reverse 5’- GTT GAA CCA GTC TCC GAC CA -3’; *Gh* forward 5’- GCT ACA GAC TCT CGG ACC TC-3’, *Gh* reverse 5’- CGG AGC ACA GCA TTA GAA AAC AG-3’; *Hgsnat* forward 5’- CGG GCG GAG CCA GAT TTA G -3’, *Hgsnat* reverse 5’- GCT CGT CCC CAA GAG TTC AT-3’; *Hprt1* forward 5’- CAG TCC CAG CGT CGT GAT TA -3’, *Hprt1* reverse 5’- GGC CTC CCA TCT CCT TCA TG -3’; *Pdia5* forward 5’- GAC CCG CAA TAA CGT GCT G -3’, *Pdia5* reverse 5’ CTC GGT CAT ACT GCA TGT GAA A -3’; *Pomc* forward 5’- ATG CCG AGA TTC TGC TAC AGT-3’, *Pomc* reverse 5’- TCC AGC GAG AGG TCG AGT TT-3’; *Scd1* forward 5’- TTC TTG CGA TAC ACT CTG GTG C -3’, *Scd1* reverse 5’- CGG GAT TGA ATG TTC TTG TCG T -3’; *sXbp1* forward 5’-GGT CTG CTG AGT CCG CAG CAG G-3’, *sXbp1* reverse 5’- AGG CTT GGT GTA TAC ATG G-3’; *Tsh* forward 5’- GGG CAA GCA GCA TCC TTT TG-3’, *Tsh* reverse 5’- GTG TCA TAC AAT ACC CAG CAC AG-3’; *Xbp1* forward 5’-AGC AGC AAG TGG TGG ATT TG -3’, *Xbp1* reverse 5’- GAG TTT TCT CCC GTA AAA GCT GA -3’.

### Chromatin immunoprecipitation assay (ChIP) assay

The chromatin immunoprecipitation assay was performed as previously reported using the SimpleChIP Enzymatic Chromatin IP Kit (Cell Signaling Technology, Cat No. 9003)^47^. Briefly, 1×10^7^ primary hepatocytes or 100 mg liver tissue were crosslinked with 1% formaldehyde (Sigma-Aldrich, Cat No. F8775), and crosslinked with 0.125 M glycine. The chromatin was immunoprecipitated with protein A/G magnetic beads (Thermo Fisher Scientific, Cat No. 88802) conjugated with anti-IgG (Cell Signaling Technology, Cat No. 2729), or anti-THRβ (Abcam, Cat No. ab196484) at 10ug/samples for 16 hrs at 4°C. The DNA was eluted from the beads and subjected to PCR analysis. The 4 sets of primers used for mouse THRβ occupancy site on *Xbp1* promoters were: set1 (−92 to −358) forward 5’- AGC CAA GGC TCT AGT GTT T -3’, reverse 5’- GCA TAT CCG GCT TAG GGT TAG -3’; set2 (−594 to −946) forward 5’- TAA GGG TGG GAC TAG GAA GAG -3’, 5’- TTA GGG AAT GGG AGT GAG GA -3’; set3 (−992 to −1228) forward 5’- CCC AAG GAG ACA TAC AGA GTC A -3’, reverse 5’- ACC TCT TTG ATA CCA CCC TAC C -3’; and set4 (−1514 to −1740) forward 5’- TTC TCT GGT GAT GAG CTG GA -3’, reverse 5’- CGG GCA TTG GTT GGC TAT ATT -3.

### Immunohistology, immunocytochemistry and electron microscopy

For immunofluorescence, frozen sections (6μm) from pituitary or liver tissues were fixed with 4% PFA, and blocked prior to overnight staining with antibodies against F4/80 (Cell Signaling, Cat No. 30325, 1:300), IRE1 alpha (Cell Signaling, Cat No. 3294, 1:300), sXBP1 (Cell Signaling, Cat No. 82914, 1:300), CHOP (Cell Signaling, Cat No. 2895, 1:300) and PERK (Cell Signaling, Cat No. 5683, 1:300), followed by 1 hr incubation with and Alexa-488-conjugated secondary antibody or Alexa-564-conjugate secondary antibody at RT for 1hr. Nuclei were stained with 4′,6-diamidino-2-phenylindole (1:2500, DAPI, ThermoFisher Scientific, Cat No. D1306). Images were taken using Zeiss 880 confocal microscopy and quantified by using Fiji/ImageJ^86^.

Human pituitary (control: BMI 21.95 and 25.6, obesity: BMI 31.1 and 44.1) samples were obtained from the Iowa NeuroBank Core, University of Iowa. The use of human pituitary tissues was approved by the institutional review board as nonhuman subject research. Sections were blocked prior to overnight staining with antibodies against IRE1α (Cell Signaling, Cat No. 3294, 1:300), XBP1 (Cell Signaling, Cat No. 12782, 1:300), followed by 1 hr incubation with and Alexa-564-conjugate secondary antibody at RT for 1 hr. Nuclei were stained with DAPI, and images were taken using Zeiss 880 confocal microscopy and quantified by using Fiji/ImageJ^86^.

To assess fibrillary collagen deposition, frozen liver tissue sections were stained by Picro Sirius Red Stain^87^. Briefly, 4% paraformaldehyde fixed sections were covered with Picro Sirius Red Solution for 60 min and washed in acetic acid. Images were observed under light microscope (Nikon).

For EM analysis, freshly isolated pituitary glands were fixed with 2.5% formaldehyde/glutaraldehyde in 0.1 M Sodium Cacodylate Buffer (pH 7.4, Emsdiasum) and 1% OsO4 (Electron Microscopy Science, Cat No. 15949), followed by dehydration and staining with uranyl acetate/lead citrate. Images were taken using a JEOL JEM 1230 electron microscope.

### Hepatic triglyceride level and Oil red O staining

Liver tissues from the same anatomical location were collected by snap freeze, and the level of hepatic TG content was determined by followed the protocol described by Triglyceride Quantification Colorimetric/Fluorometric Kit (Biovision, Cat No. K622-100) using 25 mg frozen liver sample/mouse. To measure hepatic neutral lipid, frozen liver sections (6μm) was fixed with 4% paraformaldehyde solution and stained with 0.3% Oil red O (Sigma-Aldrich, Cat No. O0625) solution. The images were taken by Nikon microscope (10X) and quantified by using Fiji/ImageJ^86^.

### Circulating hormone level and lipid profile

Serum levels of TSH (following an overnight fast), free T3 (following an overnight fast), total and free thyroxine (T4, following an overnight fast), and GH (following 6 hrs-food withdrawal) were measured using commercially available ELISA kits (TSH ELISA kit, G-Biosciences, Cat No. IT6045; free T3 ELISA kit, G-Biosciences, Cat No. IT5691; T4 ELISA kit, G-Biosciences, Cat No. IT7963; GH ELISA kit, Crystal Chem, Cat No. 80587). In addition, serum insulin levels were measured in mice after 6 hrs of food withdrawal, using a commercially available ELISA kit (ALPCO, Cat No. 80-INSMSU-E01). The plasma lipid profile, including triglyceride, cholesterol, HDL, LDL and alanine aminotransferase (ALT)/aspartate aminotransferase (AST) levels were measured in mice after 6 hrs food withdrawal using the Piccolo Lipid Panel Plus (Abaxis, Inc.)

### Whole-body metabolic profiling

Whole-body energy expenditure (VO2, VCO2), food intake, and locomotor activity were monitored using a Comprehensive Lab Animal Monitoring System (CLAMS, Columbus Instruments) at the Fraternal Order of Eagles Metabolic Phenotypic Core. The analysis of whole-body energy expenditure was used CalR (https://calrapp.org)^88^. Mouse body composition was measured by using nuclear magnetic resonance (Bruker Minispecs, LF50) at the Fraternal Order of Eagles Metabolic Phenotypic Core. Glucose tolerance was tested by measuring glucose concentration at different time points after an intraperitoneal (IP) glucose injection (1g/kg body weight, 50% dextrose, Hospra Inc, Cat No. 0409-6648-02) ^89^. Blood glucose level were determined using Bayer Contour blood glucose meter.

### Isolation of RNA and data analysis for pituitary bulk RNA sequencing

For each experimental group, total RNA was extracted from fresh liver samples or pituitary glands from 3 individual mice using Trizol and cleaned up with a RNeasy Mini Kits^47^. Quality control of isolated RNA was determined by DropSense 16 (Trinean). Pituitary RNA sequencing was performed by Iowa Institute of Human Genetics (University of Iowa) using Illumina Novaseq6000 system. The liver RNA sequencing was performed using Illumina Novaseq6000 system by Novogene. RNA-seq reads were quality checked using the FastQC tool (version, 0.1.2) and trimmed using the Java software package Trim Galore! (Version 0.6.5, Babraham Bioinformatics, Babraham Institute, Cambridge, UK). The reads were mapped to the Ensembl mouse (mm10/GRCm38) genomes using STAR software (version 2.7.10)^90^. The transcripts abundance were quantified with Salmon (version 1.8)^91^. The R Bioconductor package tximport were used to collapse our abundance estimates from the transcript level to the gene-level (version 1.22, https://f1000research.com/articles/4-1521/v2). Differentially expressed genes (DEGs) were identified by applying the R/Bioconductor package Deseq2 (version 1.34.0)^92^. Enriched pathways represented by the DEGs were identified by gene set enrichment analysis (GSEA, version 4.1.0) and Enrichr ^93^. The difference of gene between groups was based on adjusted P value <0.05.

### Pituitary scRNA sequencing

#### Pituitary gland single cell preparation

Mice were perfused via portal vein perfusion, and pituitary glands from 6-10 individual mice were dispersed with two pools of each dietary condition/genotype as previously described^94^. Briefly, the dissected glands were minced and incubated with enzyme mixture containing 2mg/ml Collagenase, Type 2 (Worthington Biochemical Corporation, Cat No. LS004176), and 5 U DNase I (Sigma, Cat No. 11284932001) at 37°C for 1hr. Cells were then washed with cold PBS, resuspended with DMEM medium (Giboco, Cat No. 10938025), and counted using hemocytometer. The cell viability was determined by trypan blue with above 90% live cells.

#### scRNA-seq library construction and sequencing

Single-cell 3′ cDNA libraries were generated using the chromium single-cell platform (V3.1) according to the manufacturer’s protocol (10× Genomics, USA), and RNA sequencing was performed by Iowa Institute of Human Genetics using Illumina Novaseq6000 system.

#### scRNA-seq data processing

FastQC (version, 0.1.2) was run with default parameters for scRNAseq QC control. For scRNAseq data analysis, Kallisto BUStool [version 0.39.1]^95^ was used to generate scRNA count matrix with the mouse indexing reference transcriptome (mm10). The scRNA seq data analysis was performed using Seurat R package (version 4.0). Data normalization was performed by “NormalizeData” in Seurat package. The cluster analysis, t-distributed stochastic neighbor embedding (tSNE), and differential gene expression analyses among the clusters were generated by Seruat package. Cells were excluded with expression genes were lower than 100, or higher than 4500^96^.

#### Cell type classification

Marker genes expressions in each cluster were identified by the function “FindAllMarkers” in Seurat package with “wilcox” methods and Bonferroni correction. The clusters were confirmed based on marker genes by the description of Cheung et al^94^. The endocrinological clusters, such as somatotrope and lactotrope, were merged by the known marker genes.

*Comparation of scRNAseq data sets and enrichment anlysis:* The different expressions of genes (DEGs) among the clusters were explored by “FindMarkers” in Seurat package. The further DEGs comparison in pro1 linage or thyrotrope clusters between RD and HFD or IRE1^fl^ and IRE1^PKO^ was used gene ontology (GO) biological process enrichment analysis of those DEGs was performed by Enrichr.

### Statistical analysis

Results are expressed as the mean ± the standard error of the mean (SEM); n represents the number of individual mice (biological replicates) or individual experiments (technical replicates) as indicated in the figure legends. Data were further analyzed with two-tailed Student’s and Welch’s t-test for two-group comparisons, ANOVA for multiple comparisons. For both One-Way ANOVA and Two-Way ANOVA, Tukey’s post-hoc multiple comparisons were applied as recommended by Prism. In all cases, GraphPad Prism (GraphPad Software Prism 8, San Diego, CA) was used for the calculations.

---

**Suppl Figure 1.**
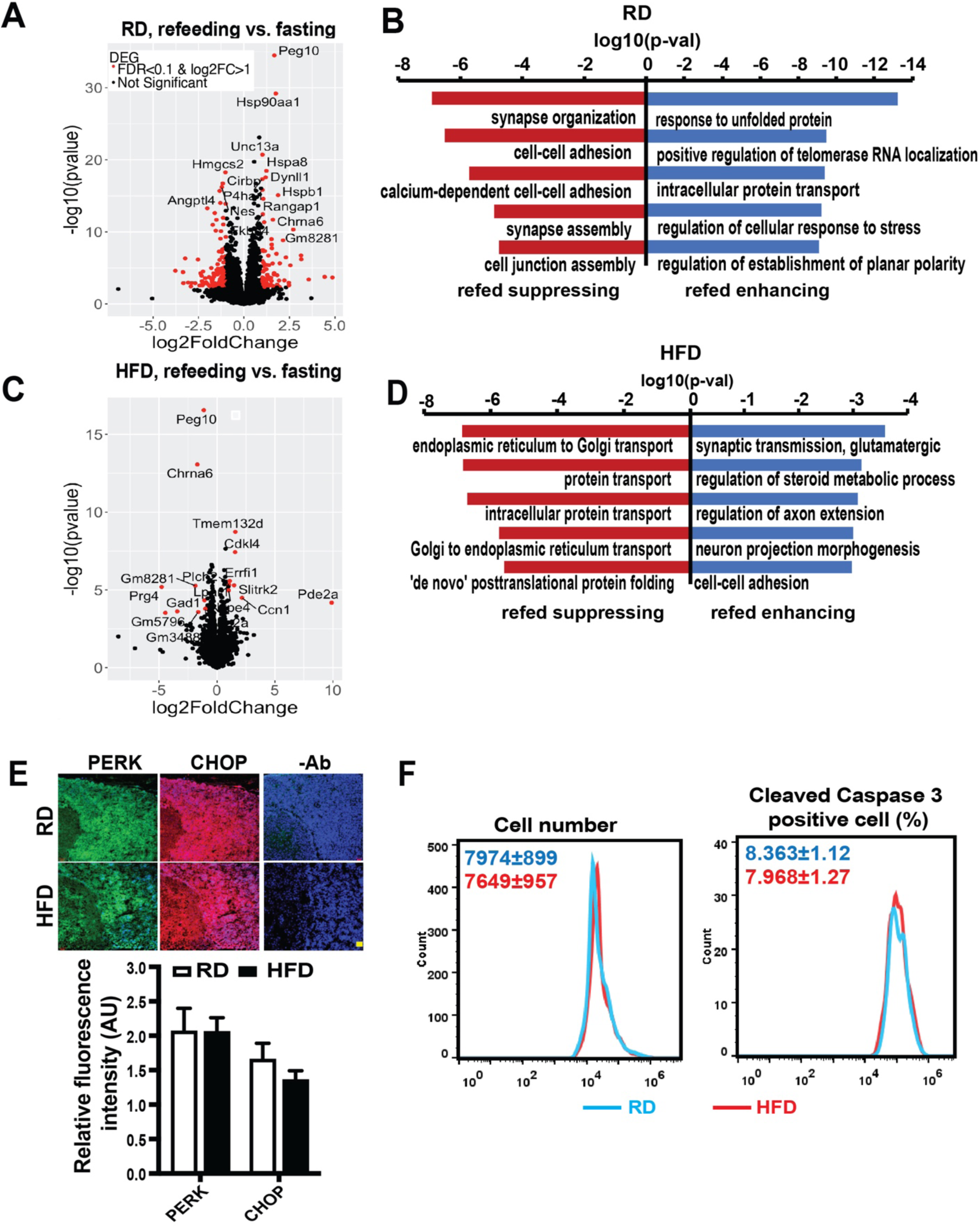
Obesity disrupted adaptive UPR in the pituitary. **A.** Volcano plot comparing RNA-seq data from pituitary of male C57BL/6J mice on a RD following a 16-hrs fasting, with or without a 4-hrs refeeding. Plot illustrates the relationship of FC (log base 2) to the p-value (−log base 10). n= 3 mice/group. **B.** Significantly enriched GO biological process pathways mice in (A) as determined by Enrichr analysis. **C.** Volcano plot comparing RNA-seq data from of pituitary of HFD mice, fasted and refed as in (A). n= 3 mice/group. **D.** Significantly enriched GO biological process pathways mice in (C) as determined by Enrichr analysis. **E.** Representative images and quantification of tested proteins in the pituitary from RD and HFD (16 wks on a HFD) male mice following a 16-hrs fasting with a 4-hrs refeeding, n= 4 mice/group. Scale bar: 50 μm. **F.** Flow cytometry analysis in anterior pituitary cells from mice in C57BL/6J mice fed a RD and HFD for 16 wks. n= 3 mice/group. In E and F, data are presented as means ± SEM and statistical analysis was performed by Student’s t-test.

**Suppl Figure 2.**
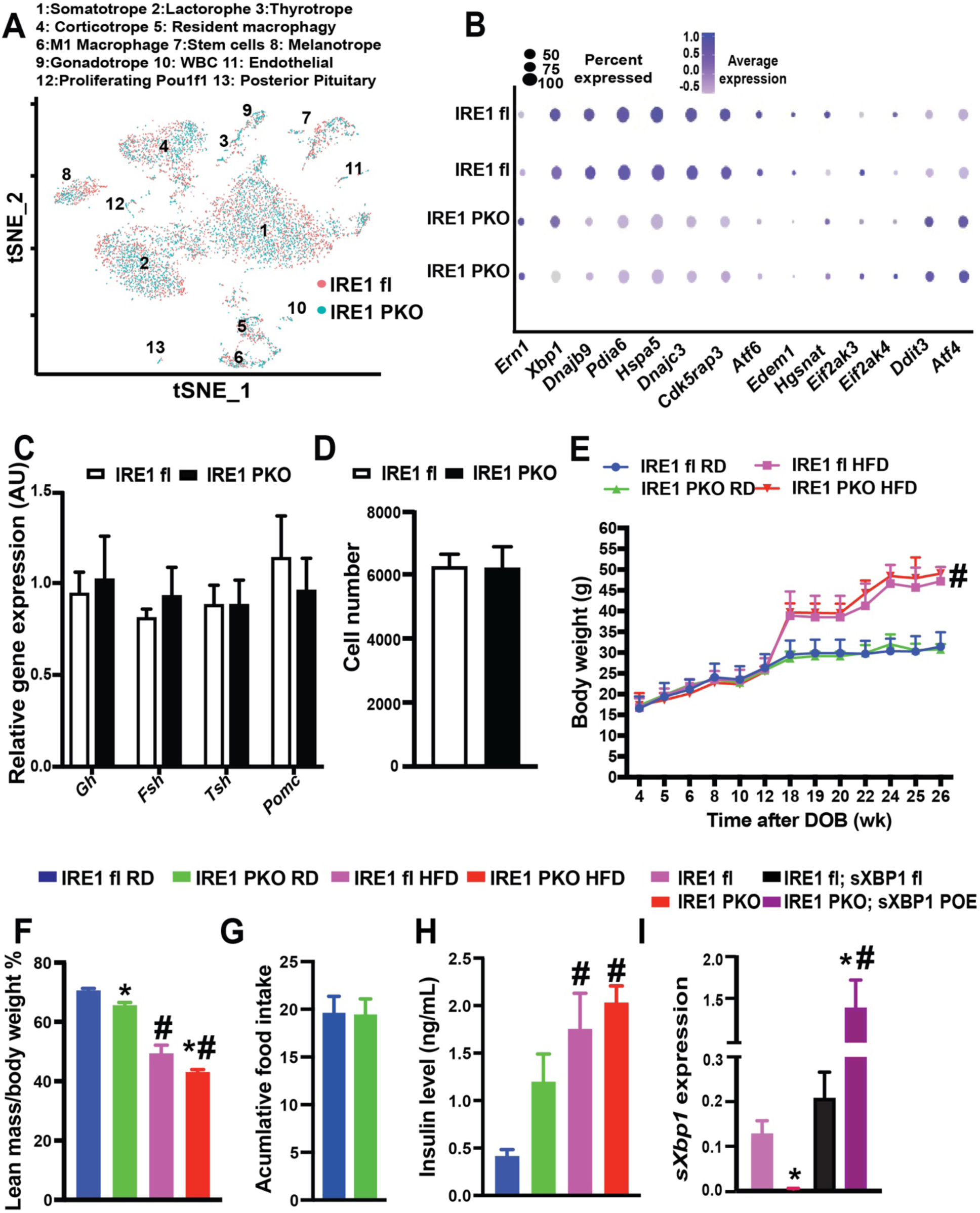
Metabolic profiles of IRE1α pituitary deficient mice. **A.** tSNE plot and **B.** Dot plots of thyrotrope-UPR markers in pituitary of IRE1^fl^ and IRE1^PKO^ mice on a RD. **C.** Levels of mRNAs encoding genes of interest in pituitary mice in (A), as assessed by quantitative RT-PCR. n= 3 mice/group **D.** Cell numbers of anterior pituitary isolated from mice in (A) measured by flow cytometry analysis. n= 3-4 mice/group**. E.** Body weight, **F.** lean body mass, **G.** food intake, **H**. circulating insulin level in IRE1^fl^ and IRE1^PKO^ mice fed a RD or a HFD. n= 5-6 mice/group**. I.** *sXbp1* expression in pituitary from IRE1^fl^, IRE1^PKO^, IRE1^fl^; sXBP1^fl^ and IRE1^PKO^; sXBP1^POE^ mice fed on a HFD for 20 wks. n= 3 mice/group. Data are presented as means ± SEM in (C-I). * Indicates genetic effects in mice with same diet, and # indicates dietary effects in same type of mice; as determined by Student’s t-test in (C, D and G), and ANOVA followed by post-hoc test in (E, F, H and I), p<0.05.

**Suppl Figure 3.**
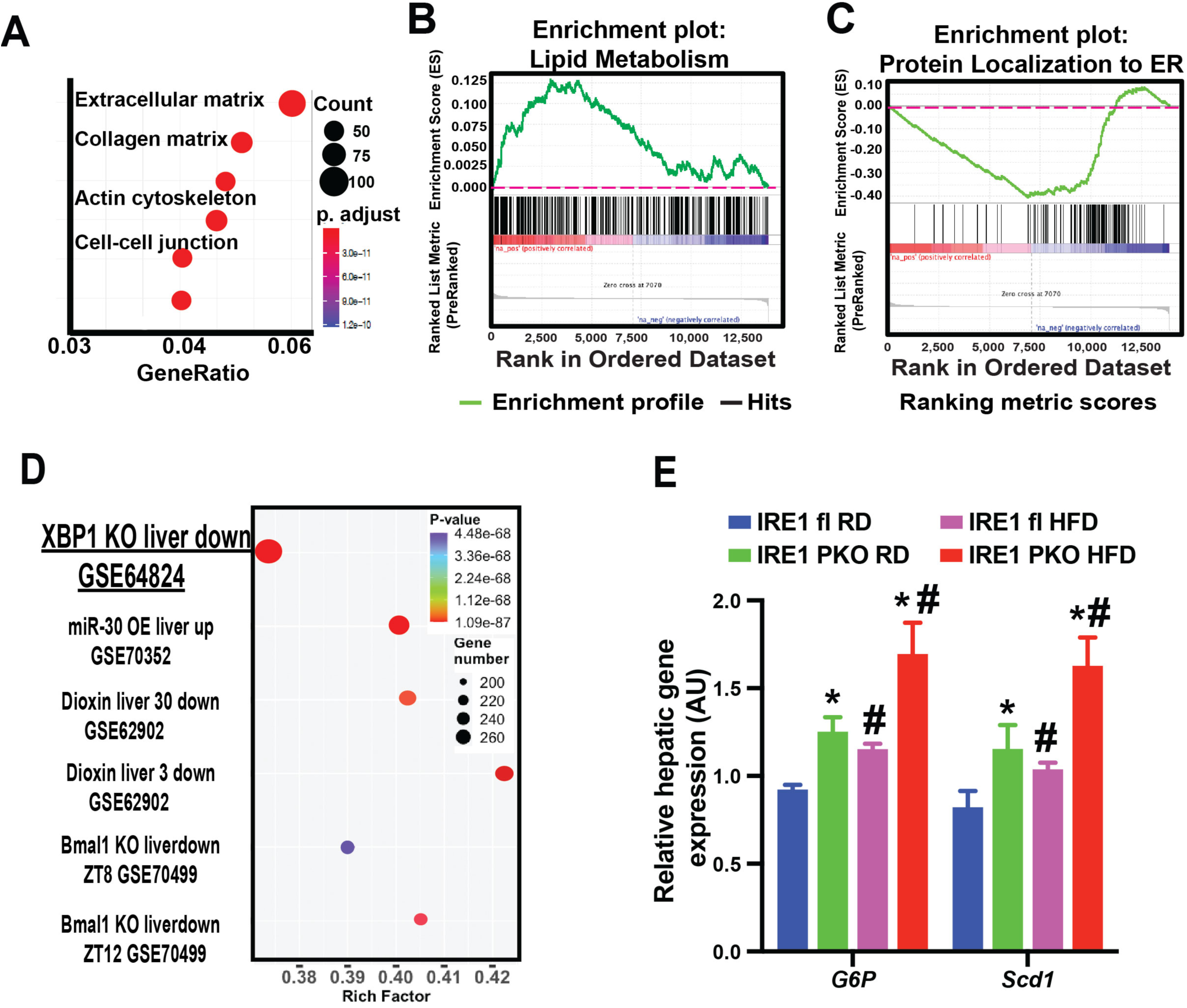
The effects of pituitary IRE1α deletion on hepatic metabolic function. **A.** GO analysis of up-regulated DEGs in hepatic RNAseq analysis in IRE1^fl^ and IRE1^PKO^ mice on HFD for 20 wks. n= 3 mice/group. **B.** GSEA analysis of DEGs of lipid metabolism, and **C.** protein localization to ER in liver as described in (A). **D.** RNA-Seq Disease Gene signatures of livers from IRE1^fl^ and IRE1^PKO^ mice on a HFD for 20 wks. **E.** Levels of mRNAs encoding genes of interest in livers from mice in (A) as assessed by quantitative RT-PCR. Data are presented as means ± SEM in (E). * Indicates genetic effects in mice with same diet, and # indicates dietary effects in same type of mice as determined by ANOVA followed by post-hoc test, p<0.05.

**Suppl Figure 4.**
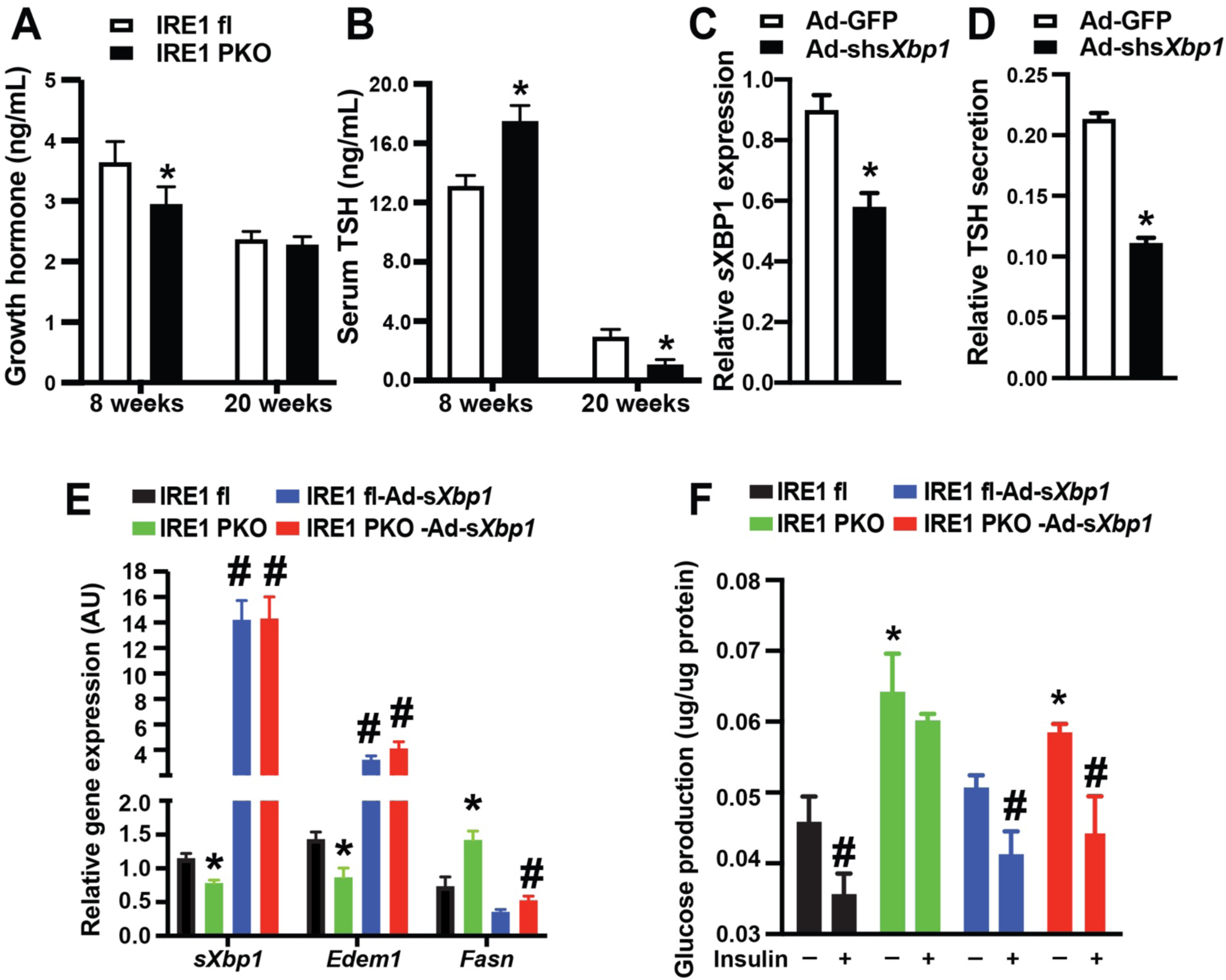
The role of pituitary UPR-TH-liver axis on the progression of NAFLD. **A&B.** Serum levels of hormone of interest in IRE1^fl^ and IRE1^PKO^ mice on a RD at age of 8 and 20 wks. n= 4-6 mice/group **C.** sXBP1 expression in isolated pituitary from WT mice measured by indirect ELISA. Pituitary glands were transduced with indicated adenovirus at a titer of 2×10^6^ VP/ml for 48 hrs. n= 3 mice/group**. D.** TRH-stimulated TSH levels in anterior pituitaries as in (C). **E**. Level of mRNAs encoding genes of interest in liver from IRE1^fl^, IRE1^PKO^ mice transduced with adeno-s*Xbp1* on a HFD for 16 wks. **F.** Hepatic glucose production in primary hepatocytes isolated from mice in (E). Cells were treated with 100 nM insulin and 10 µM forskolin. n= 3 mice/group. All data are presented as means ± SEM. * Indicates genetic effects in cells or mice with same treatment in (A-D), and # indicates treatment effects in same type of mice as determined by ANOVA and followed by post-hoc test in (E&F), p<0.05. AU: arbitrary unit.

